# Revisiting the interaction of heme with hemopexin: Recommendations for the responsible use of an emerging drug

**DOI:** 10.1101/2020.04.16.044321

**Authors:** Milena S. Detzel, Benjamin F. Syllwasschy, Francèl Steinbock, Anuradha Ramoji, Marie-Thérèse Hopp, Ajay A. Paul George, Ute Neugebauer, Diana Imhof

**Author notes:** M.S.D. and B.F.S. contributed equally to this work. Correspondence and requests for materials should be addressed to D.I.

## Abstract

In hemolytic disorders, erythrocyte lysis results in massive release of hemoglobin and, subsequently, toxic heme. Hemopexin is the major protective factor against heme toxicity in human blood and currently considered for therapeutic use. It has been widely accepted that hemopexin binds heme with extraordinarily high affinity in a 1:1 ratio. Here we show that hemopexin binds heme with lower affinity than previously assumed and that the interaction ratio tends to 2:1 (heme:hemopexin) or above. The heme-binding sites of hemopexin were characterized using hemopexin-derived peptide models and competitive displacement assays. In addition, *in silico* molecular modelling with a newly created homology model of human hemopexin allowed us to propose a recruiting mechanism by which heme consecutively binds to several histidine residues and is finally funnelled into the high-affinity binding pocket. Our findings have direct implications for the biomedical application of hemopexin and its potential administration in hemolytic disorders.

## Main Text

### Introduction

In hemolytic conditions, occurring e.g., in malaria, sickle cell disease, and thalassemia, erythrocytes undergo lysis and large amounts of hemoglobin (≥20 μM) enter the blood stream^1–4^. In healthy patients, released hemoglobin is directly bound by haptoglobin and transported to hepatocytes and macrophages for its degradation^5^. This protective mechanism, however, can collapse when the scavenging capacity of haptoglobin is exhausted^4^. Then, hemoglobin is oxidized to methemoglobin, and heme (Fe(III) protoporphyrin IX) is released into the blood^4^. Serum albumin was suggested to unspecifically bind heme and to transfer it to the major physiological heme scavenger hemopexin^5,6^. Hemopexin binds heme specifically with a serum capacity of approximately 17 μM^7^ and escorts it to the liver, where the complex is subsequently degraded^8^. Patients faced with elevated heme levels often develop severe complications, including stroke, deep venous thrombosis, and pulmonary embolism^3,9^. With increasing concentrations, labile heme can generate dangerous reactive oxygen species (ROS) by participating in Fenton-type reactions with hydrogen peroxide as substrate^10^. As a consequence of ROS production, oxidative damage may occur to proteins, carbohydrates, lipids, and nucleic acids^4^. In addition, heme itself can act as regulator by transiently binding to proteins^11,12^. It can directly alter the function of several plasma proteins that are part of the immune system e.g., IgG, C1q, and TLR4, and exacerbate other pathologies^13–15^. Consequences of the accumulation of labile heme are hepatic injuries in sepsis, nephropathy in hemolysis patients, and leukocyte and reticulocyte adhesion to endothelial cells^16–18^. The latter may be responsible for vascular occlusion in patients with sickle cell disease^17^. Hemopexin knockout mice are viable, but showed neuronal degeneration, striatal injuries and lethargy after artificially induced intracerebral hemorrhage^19^. Consequently, researchers sought to apply hemopexin as therapeutic agent when the bodies’ heme-scavenging capacity is exhausted. There are abundant model systems of human diseases, in which hemopexin administration improved clinical parameters, which have been reviewed elsewhere^20^. These promising results currently fuel commercial interest in plasma-derived hemopexin as potential treatment for e.g., sickle cell disease.

The nature of the heme-hemopexin complex was intensively investigated from 1970 to 2000, when researchers were interested in the K_D_ value, ratio, and binding affinity of this interaction^21^–^26^. The first dissociation constant (K_D_ <10 nM) with a 1:1 (heme-hemopexin) ratio for heme binding to rabbit hemopexin was determined via photometric titrations in 1972^24^. The method, however, suffered intrinsic detection limitations^24^ and, thus, the transfer of ferrihemoglobin-bound heme to human hemopexin was used in an alternative assay in 1974^26^. Hemopexin was able to acquire heme from the isolated hemoglobin alpha chain and a K_D_ of <1pM was derived based on the known association constant of heme and globin (0.19 pM)^27^. Today, sophisticated biophysical methods for direct analysis and real-time monitoring of biomolecular interactions are available^28^. Almost 50 years after the introduction of binding constants for the heme-hemopexin interaction, it seems appropriate to inspire and promote reflection and a debate of the former analyses.

For understanding the hemopexin function in physiology and pathology and its subsequent therapeutic use, it is also important to know the exact heme-binding capacity of hemopexin. Several conserved histidines, especially H32, H79, H150, and H236 (numbering as in the human protein), were initially suggested as possible heme-coordinating sites^21,23,29,30^. NMR and CD spectroscopy also supported the notion of multiple heme-binding sites in hemopexin^22,31^. Notably, the aforementioned residues are conserved in rat, rabbit, and human hemopexin^30^. H79 and H150 were suggested as heme-binding sites by using chemically modified hemopexin domain I isolated from rabbit serum^21^, since they were protected against carboxymethylation after complex formation with heme. Participation of H32 and H236 was excluded in this study, because no change in absorbance was found after their removal upon proteolytic cleavage^21^. Recombinantly expressed hemopexin double mutant H79T/H150T, but not H79T alone, revealed a ten-times lower heme-binding affinity compared to wild-type hemopexin, which suggested a supporting role of H79^29^. Surprisingly, in the first crystal structure of heme-bound rabbit hemopexin, the binding sites were shown to be H236 and H293, which complexed one heme molecule in a hexacoordinate state^23^. In the same work, H105 and H150 were suggested as possible additional binding sites^23^.

The discrepancy between the coordinating histidine residues of the crystal structure and the ones suggested in earlier titration experiments prompted us to explore heme binding to all available heme-binding motifs (HBMs)^23,29^ of hemopexin and to re-evaluate the earlier reports. Our data reveal that the K_D_ value for the heme-hemopexin interaction is in the subnanomolar range and the stoichiometry exceeds the 1:1 ratio suggested earlier. In addition, we generated a homology model of human hemopexin to facilitate molecular docking and molecular dynamics simulations to gain insight into the underlying process of heme binding to hemopexin, which support a newly defined recruiting mechanism and more than one heme-binding site in hemopexin.

## Results

### Heme binds to human hemopexin in the subnanomolar range

To characterize the binding of heme to hemopexin with the aim to re-examine the published K_D_ values (<10 nM^24^ and <1 pM^26^, respectively), SPR analysis was performed as described earlier for other heme-binding proteins (Fig. 1a)^11^. Experimental data of the first, high-affinity heme-hemopexin interaction was fitted to a global 1:1 kinetic model (k_a_ = 1.14 ± 0.24 x 10^6^ M^-1^s^-1^, k_d_ = 3.61 ± 0.42 x 10^-4^ s^-1^) yielding a K_D_ value of 0.32 ± 0.04 nM. This value is higher than the suggested <1 pM from Hrkal *et al*. in 1974^26^, but lower than the estimated value of 10 nM reported by Morgan *et al*. in 1972^24^. The binding stoichiometry for this initial binding event at heme concentrations from 0.75 nM to 12 nM was determined to be approximately 1:1 (heme:hemopexin). At higher heme concentrations (>100 nM) the stoichiometry changed to >2:1. However, a strong unspecific signal increase, which might be due to heme aggregation effects, prevented further SPR analysis of the second heme-binding event. The SPR experiments indicated that more than one heme-binding site might exist in hemopexin. To confirm this finding, we performed a titration experiment of hemopexin with heme using UV/Vis spectroscopy and observed an increase of the Soret band at 415 nm even at ratios exceeding 2:1 (heme:hemopexin) (Supplementary Fig. 1). However, in contrast to the previously described heme-IL-36α interaction, only one maximum was observed in the spectra, which prevents a separate evaluation of independent maxima, such as for the CP- and HY-binding site as previously shown in IL-36α^11^. All reported heme-binding motifs of hemopexin contain histidine as heme-coordinating amino acid (*vide infra*). Consequently, UV/Vis shifts of all histidine-based heme-binding sites overlap and form one maximum. A nonlinear fit of the data^32^ revealed a stoichiometry of 2:1 (heme:hemopexin) for this interaction and a K_D_ value of 3.53 ± 0.80 μM, which represents a mixed value for all binding sites. These results substantiate the binding behaviour observed in SPR, as well as that of previous reports, in which the isolated N-terminal hemopexin domain, which lacks the confirmed binding site around H236/H293, binds heme^21,23^. Furthermore, the combined results suggest that hemopexin harbours one high- and at least one low-affinity heme-binding site.

**Fig. 1.**
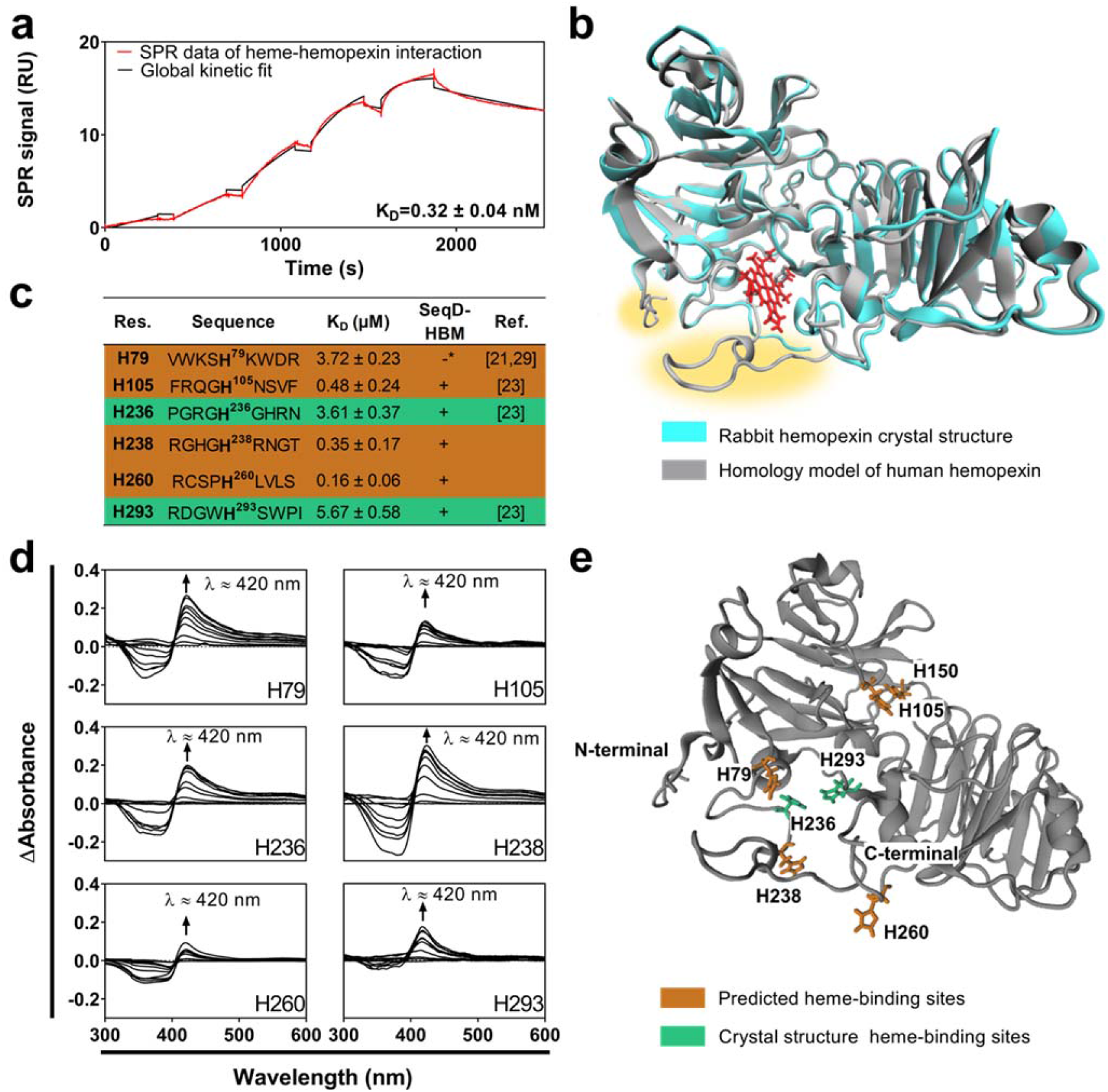
Hemopexin binds heme with lower affinity than assumed and contains additional surface-exposed heme-binding sites. **a** SPR signal (RU) of hemopexin with increasing heme concentrations (0.75 nM to 12 nM) is shown in red, the single-cycle kinetics fit is displayed in black. Experiments indicated a stoichiometry exceeding 2:1 (heme:hemopexin). **b** Homology model of human hemopexin, shown in grey ribbons compared to the crystal structure of rabbit hemopexin (PDB ID:1QJS)^23^ shown in cyan ribbons with heme shown in red sticks in the known binding pocket involving H236 and H293. The structural alignment between the crystal structure and the homology model resulted in a backbone RMSD of 0.935 Å over 396 aligned residues with 84.34% sequence identity. The 15.66% difference in sequence identity is mostly accounted for differences in the region of residues 2 to 5 and 242 to 256, which form the flexible hinge loop (highlighted in yellow). **c** The algorithm SeqD-HBM predicts further heme-binding sites. Residue number (Res.), sequences of the synthesized peptides, determined K_D_ values respective reference (Ref.). The two motifs shown in the crystal structure^23^ are highlighted in green, while predicted heme-binding sites are marked in orange. *This sequence was excluded by WESA. d Difference spectra (spectrum of pure heme subtracted from spectrum of heme-peptide complex) of motifs H79, H105, H236, H238, H260, H293 binding to heme. All complexes show a shift of the Soret band to around 420 nm (marked by an arrow), which indicates heme binding^36^. e Visualization of known (green) and predicted (orange) heme-binding motifs in the homology model. All motifs, except for H105 and H150, are in close proximity to the known heme-binding site and are surface accessible.

### *In silico* homology modelling of hemopexin

In order to enable computational studies, homology modelling of hemopexin was performed, as no structure for the human wild-type protein is available. YASARA version 19.9.17^33^ was used to generate the homology model based on the X-ray analysis of the rabbit protein (398 amino acids, PDB-ID: 1QJS) and addition of the missing residues 99-105 and 216-221 of the human protein^23^. The homology model achieved an optimal Z-score of −1.068, which reflects an excellent alignment with the available structure^23^ (Fig. 1b). The structural alignment between the crystal structure and the homology model resulted in a root-mean-square deviation (RMSD) value of 0.935 Å over 396 aligned residues with 84.34% sequence identity between human and rabbit hemopexin. The 15.66% difference in sequence identity is mostly accounted for differences in the region of residues 2 to 5 and 242 to 256, which form the flexible hinge loop.

### Analysis of potential heme-binding motifs in hemopexin

Considering that one heme-binding site in the crystal structure consists of residues H236 and H293^23^, the question arises where additional binding site(s) could be located. At least three additional histidine residues have been suggested as heme-coordinating sites in earlier studies, namely H79, H105, and H150^23,29^. We used the recently introduced heme-binding motif (HBM) search algorithm SeqD-HBM^34^ to screen the hemopexin sequence for all potential heme-binding sites. SeqD-HBM identified 11 nonapeptide motifs in hemopexin, all based on coordination by histidine (8 motifs) or tyrosine (2 motifs) (Supplementary Tab. 1). The earlier suggested H150^23,29^ was excluded by the algorithm due to an intramolecular disulfide bond occurring within the motif AVE**C**HRGE**C**. In addition, H79 was not detected by the algorithm due to an insufficient solvent (i.e. surface) exposure as was determined by the WESA (*Weighted Ensemble Solvent Accessibility*) module^34^. However, because of their occurrence in previous publications, we added these two motifs to the predicted sequences and synthesized 13 nonapeptide motifs for in-depth analysis of their heme binding. The motifs H150, C154, Y254 and H260 (Supplementary Tab. 1) contain cysteine residues, which may be present in the oxidized form in the protein. In order to clarify the oxidation state of these cysteines, we incubated hemopexin with the alkylating agent 2-iodoacetamide and performed mass spectrometry measurements^35^. This analysis revealed that all cysteines are oxidized in wild-type hemopexin, which was also confirmed by a negative result from the Ellman’s test for free thiols^35^. Thus, peptide H150 was synthesized with an intramolecular disulfide bond and in all other peptides, disulfide bonds were mimicked by insertion of a methylated cysteine during synthesis.

### Heme binds to hemopexin-derived peptides

The heme-binding behaviour of the peptides was analysed by UV/Vis spectroscopy as established previously^36^. Six out of 13 peptides, namely H79, H105, H236, H238, H260, and H293, showed a bathochromic shift of the Soret band to around 420 nm indicating heme binding (Fig. 1c, d). Subsequently, K_D_ values for the heme-peptide interactions were determined from the difference spectra (Fig. 1d). The peptide possessing the highest heme-binding affinity was H260 (RCSP**H**LVLS, K_D_ = 0.16 ± 0.06 μM). The peptide motif derived from H105^23^ showed a high affinity for heme (FRWG**H**NSVF, K_D_ = 0.48 ± 0.24 μM), as well, which can be explained by the presence of hydrophobic aliphatic and aromatic (V, F, W) as well as positively charged (R, H) residues^34,37^. Motifs H236 (PGRG**H**GHRN, K_D_ = 3.61 ± 0.37 μM) and H293 (RDGW**H**SWPI, K_D_ = 5.67 ± 0.58 μM), i.e. the two heme-coordinating sites in the crystal structure of rabbit hemopexin^23^, showed only moderate heme-binding affinity. Here, efficient heme binding might be due to the formation of a specific heme-binding pocket in the protein context^23^, which is mediated by a large conformational change upon heme association^38^. The heme-binding amino acids H236 and H238 are contained in two peptides of this study. Indeed, these residues might be considered a HXH motif^32^. While peptide H236 was a moderate binder, the motif H238 displayed high heme-binding affinity (RGHG**H**RNGT, K_D_ = 0.35 ± 0.17 μM). The sequence surrounding H79 (VWKS**H**KWDR), which was earlier suggested as potential heme-binding site^21,29^, also displayed only moderate affinity (K_D_ = 3.72 ± 0.23 μM) compared to H105, H238 and H260. Unfortunately, structural insights into the heme coordination states could not be obtained so far, because the solubility of these heme-peptide complexes was too low for 2D NMR analysis. We thus performed resonance Raman (rRaman) spectroscopy measurements (Supplementary Fig. 2) and were able to acquire spectra for the heme-peptide complexes of motifs H79, H105, H236, H260, H293, and H150, while the heme complex of H238 was not sufficiently soluble. All examined peptides bound heme in a pentacoordinate state as indicated by a clear *ν*3 vibrational band^36^, with the exception of peptide H293, which showed a mixture of penta- and hexacoordination for its heme complex^36^. In the context of the full-length protein, this situation could be realized by hexacoordinate heme binding between H236 and H293 as occurring in the crystal structure^23^. As the surface-accessibility of an HBM is important for potential heme binding to the model, the predicted heme-binding sites were visualized (Fig. 1e) and all were deemed to be surface-accessible.

### Transfer of heme from peptides and albumin to hemopexin

*In vitro* and in the human body, heme is transferred from albumin to hemopexin^5,6,39^. We hypothesized that there could be an internal transfer of heme involving different motifs of hemopexin as part of a recruiting mechanism. Consequently, we investigated the differences regarding the transfer of heme from several of the aforementioned heme-peptide complexes to hemopexin.

The complexes of heme with the motifs H105, H236, H238, H260, and H293 were formed in a 1:1 ratio. The motifs H238 and H260 were chosen because of their high heme-binding affinity, whereas H105 was earlier suggested as heme-binding motif^23^ and shown to bind heme with high affinity in the UV/Vis studies. The motifs represented by the peptides H236 and H293 were determined as heme-binding sites in the crystal structure analysis^23^. Preformed heme-peptide complexes were incubated with hemopexin in a 2:1 ratio (complex:hemopexin) (Fig. 2a). The transfer of heme from the individual heme-peptide complex to hemopexin was observed as a time-dependent shift of the Soret band over 40 min (Fig. 2b). After 40 minutes, the absorbance of the Soret band was higher than that of hemopexin incubated with equimolar amounts of heme (Supplementary Fig. 1). From this observation it was concluded that hemopexin was able to accept at least one heme molecule from the heme-peptide complexes. The same experiment was also performed with human serum albumin (HSA) as previously reported by others^5,6,39^. With this experiment, we were able to confirm the results observed by Morgan and co-workers in 1976, i.e. hemopexin is able to extract at least one heme molecule from HSA^6^. Comparing the peptides with HSA revealed differences in the velocity of the heme transfer as follows: HSA > H260 > H293 > H236/H238 > H105 (Fig. 2c). These data suggest that a transfer of heme between several heme-binding sites in hemopexin is possible in the protein context as well. The difference between heme-transfer velocities supports the notion that hemopexin can accept heme more readily when it is already bound by another peptide or protein. This behaviour points towards a possible intramolecular recruiting mechanism. Interestingly, transfer from HSA to hemopexin was faster than from hemopexin-derived nonapeptides, which is in line with the physiological situation, in which this transfer is crucial.

**Fig. 2.**
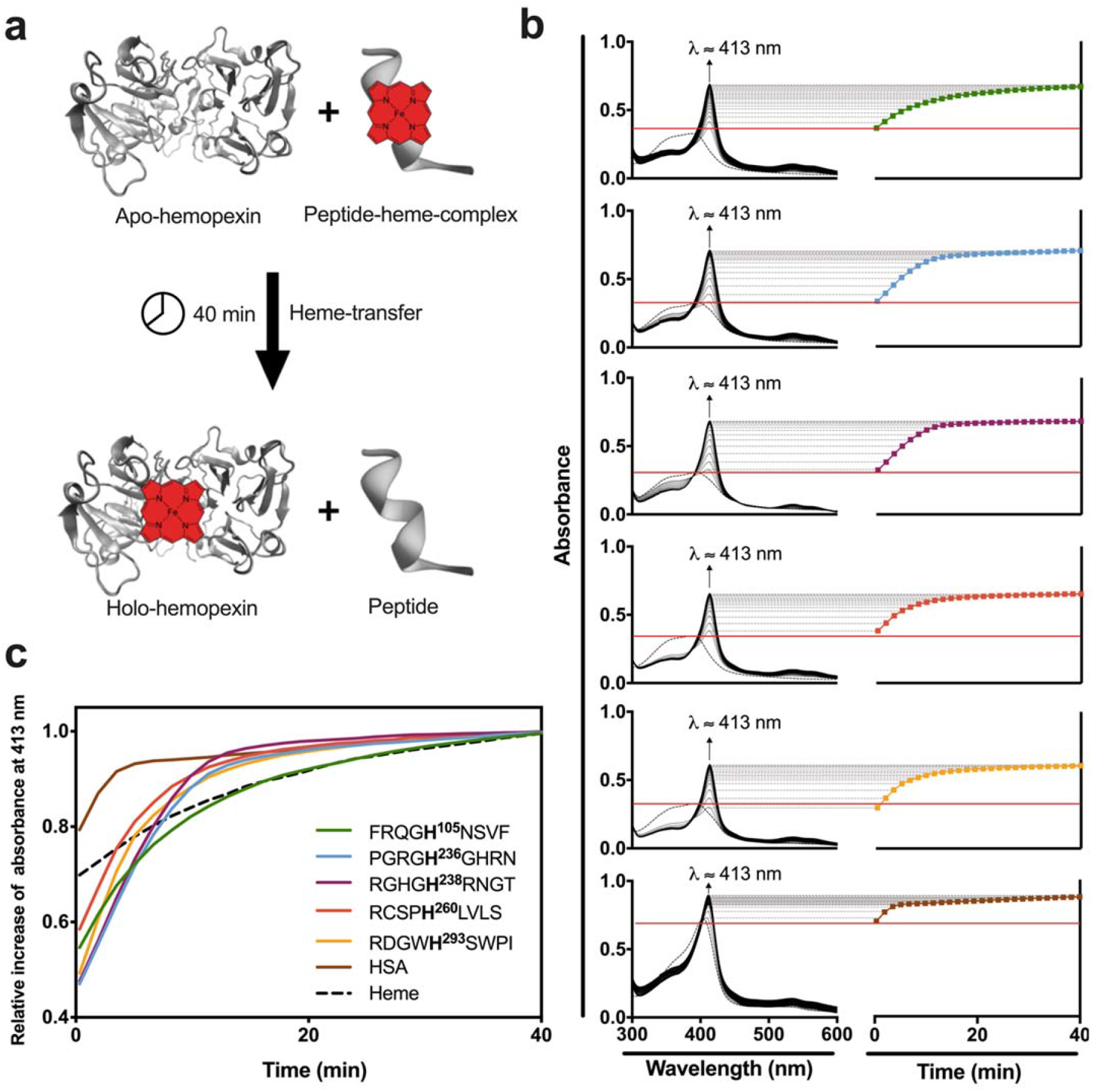
Transfer of heme from heme-peptide/HSA complexes to hemopexin. **a** Hemopexin (5 μM) was added to a heme-peptide/HSA-complex solution (10 μM), resulting in a 1:2 (hemopexin:complex) ratio, and the absorbance increase of the heme transfer was monitored for 40 minutes. **b** The resulting raw spectra for each heme-peptide complex show a time-dependent increase of the Soret band at 413 nm, which is typical for the heme-hemopexin complex. The dotted black line represents the spectrum of the respective heme-peptide/HSA-complex before addition of hemopexin. The red line highlights the maximum of the Soret band of the heme-peptide/HSA-complex before adding hemopexin. **c** Comparison of the relative increase in Soret band intensity, which represents the velocity of heme transfer. Transfer velocities decrease in the order HSA (solid black line) > H260 > **H293** > **H236**/H238 > H105 (bold residues are the confirmed heme-binding sites in the crystal structure^23^).

### *In silico* study of hemopexin by molecular dynamics simulations

To gain insight into the existence of a possible heme-recruiting mechanism, we examined both holo-hemopexin (containing heme) and apo-hemopexin (without heme) in this study. The homology model, which initially contained heme, was directly used as the holo-hemopexin model. To generate the apo-hemopexin model, heme was removed from the homology model. MD simulations were conducted to generate a conformational ensemble of the protein models to observe their individual structural dynamics and to ensure that the structures were stable. Both apo-hemopexin and holo-hemopexin were subjected to 200 ns molecular dynamics (MD) simulations using the AMBER11 force field^40^. MD trajectory analysis based on the evolution of the backbone RMSD profiles indicated that both models were stable after their simulations, with structural stability reached at around 175 ns (Supplementary Fig. 3). The observed structural dynamics were similar for both the apo- and holo-hemopexin, with the greatest structural fluctuations being in the N-terminal region and in the regions of residues 230 to 260, which is part of the *hinge* region. This region is of particular interest, as it is part of the known heme-binding pocket^23^. The fluctuations in this region give rise to a flexible loop between the two domains of hemopexin, which is found in both the apo- and holo-hemopexin and seems to play a role in the binding of heme to hemopexin. The analysed results were indicative of stable protein structures and the apo- and holo-hemopexin models were deemed suitable structures to further investigate the possibility of a heme-recruiting mechanism and a heme-protein binding ratio exceeding 1:1.

### *In silico* docking of heme to apo- and holo-hemopexin

The mechanisms of heme binding and the experimentally suggested higher heme-hemopexin binding ratio were investigated by molecular docking. Heme was successfully docked to the known H236/H293 binding site^23^ in apo-hemopexin (Supplementary Fig. 4). Subsequently, docking of heme to the apo-hemopexin model was successful with all the experimentally determined HBMs forming pentacoordinate bonds between the histidine nitrogen (N_ε_) atom and the heme iron ion. Heme docking to the predicted HBMs on the apo-hemopexin structure confirmed the experimental results that motifs around H79, H238 and H260 are good binders (Supplementary Fig. 5).

The heme-H79 docked complex gave the lowest binding energy and an in-pocket binding complex. The orientation of H79 (Fig. 3a) facilitates the orientation of heme into the binding pocket determined in the crystal structure^23^. A hydrogen bond between N246 and heme stabilizes the binding, as well as the unique hydrophobic, aromatic composition of the binding pocket (W194, Y199, F206, Y220, Y227, F228, W295)^23^.

**Fig. 3.**
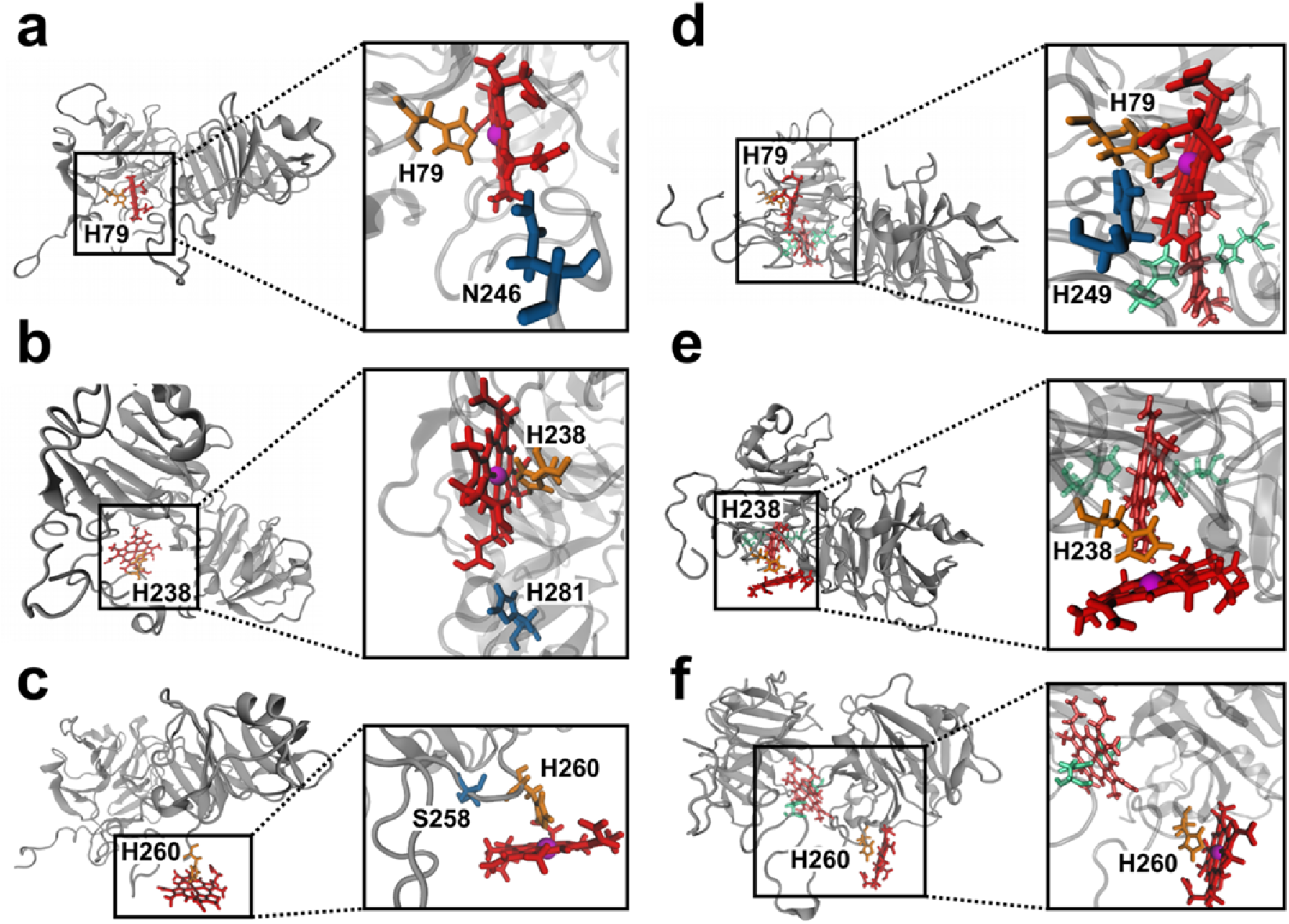
Heme docking to predicted motifs in apo- and holo-hemopexin. Heme docks to three previously predicted heme-binding sites through histidine coordination of the iron ion. Overview and close up of the apo- (left) and holo-hemopexin (right) structures (grey ribbons), binding one heme (red sticks) via predicted histidines (orange sticks) and supporting amino acids (blue sticks). The holo-hemopexin structure is depicted with the first heme (bright red sticks) in the known pocket bound by H236 and H293 (green sticks). **a** H79 is located close to the crystal structure’s binding pocket. A hydrogen bond between N246 (H atom) and one heme propionic acid (O atom) acts as a supporting bond. **b** H238 is situated in the crystal structure’s binding pocket^23^. A hydrogen bond between H281 (H atom) and one heme propionic acid (O atom) act as a supporting bond. **c** H260 is situated on the outer surface of the protein pointing outwardly. A hydrogen bond between S258 bond (H atom) and one heme propionic acid (O atom) acts as a supporting bond. **d** With docking a second heme it can be seen that heme is able to bind to H79 inside the crystal structure’s binding pocket. A hydrogen bond between H249 (N atom) and heme (H atom) acts as a supporting bond. **e** H238 is pointed outwardly to bind heme in the docking of a second heme close to the binding pocket. **f** A second heme is able to bind to hemopexin via H260 with some distance to the binding pocket.

Similar to H79, heme formed an in-pocket binding complex in case of the heme-H238 association with two stabilizing hydrogen bonds formed with R239 and H281, respectively (Fig. 3b)^23^. In the complex of heme and motif H260, heme is orientated away from the protein’s surface and therefore binding of heme occurs on the surface of the apo-hemopexin structure (Fig. 3c). The binding complex is supported by a hydrogen bond between S258 and heme. Although both H105 and H150 were able to bind heme in the docking experiment, a binding energy analysis indicated that in comparison with the other HBM’s, H105 and H150 are weaker binders (Supplementary Fig. 5). The weaker binding capability can be explained by the positioning and the suboptimal solvent-accessible area of the two motifs. These two motifs were neither fully exposed as in the case of H260 nor were they involved in in-pocket binding as H238. The described docked complexes remained stable during the 500 ps refinement simulation^41^, which was used to validate the complexes in solution, as well as during the two 20 ns MD simulations.

To gain more insight into the suggested higher heme-hemopexin binding ratio, docking was also conducted on the holo-hemopexin model. The docking of heme to the experimentally determined HBMs on the holo-hemopexin structure were successful with all the motifs being able to form pentacoordinate complexes via a bond between the histidine nitrogen (N_ε_) atom and the heme iron ion. The positioning and orientation of the H79 in the known binding pocket made it the best binding HBM in the case of holo-hemopexin (Supplementary Fig. 5). Interestingly, the binding pocket was able to accommodate the additionally introduced heme. The binding of heme to H79 (Fig. 3d) was stabilized by a hydrogen bond formed between H249 and the binding complex. The binding energy of all the other heme-HBM docked complexes were similar, with H105 showing the weakest binding energy (Supplementary Fig. 5). The weak binding of H105 can be attributed to poor surface-accessibility and the distance from the preferred binding site. As the binding energies are only approximations, the calculated binding energies for H150, H238, and H260 were too similar to distinguish between their binding capabilities (Supplementary Fig. 5). Closer visual inspection of the docked complexes indicated that H238 (Fig. 3e) and H260 (Fig. 3f) may be more likely to form binding complexes than H150, as they are more exposed to the surface. The docked complexes remained stable over the course of the 500 ps refinement simulation, as well as during the two 20 ns MD simulations, indicating that hemopexin may be able to bind more than one heme molecule at a time.

### Mechanism of heme recruitment by hemopexin

From the molecular docking and MD simulations, it seems probable that a recruitment mechanism may exist. Possible mechanisms are: (i) H260 initially binds heme on the surface of the protein with H236 and H293 already in their correct positions for binding, movement in the flexible loop causes H260 to shift into a less favourable position. This shift may propagate the transfer of heme from H260 into the binding pocket (Fig. 4a). In the other scenario (ii) H79 binds heme in the known binding pocket with H293 already in position. H236 is, however, not yet in a favourable position to form the H236/H293 binding complex. Movements in the flexible loop, relocate H236 into an ideal binding position, with the positioning of H79 becoming unfavourable for heme binding (Fig. 4b).

**Fig. 4.**
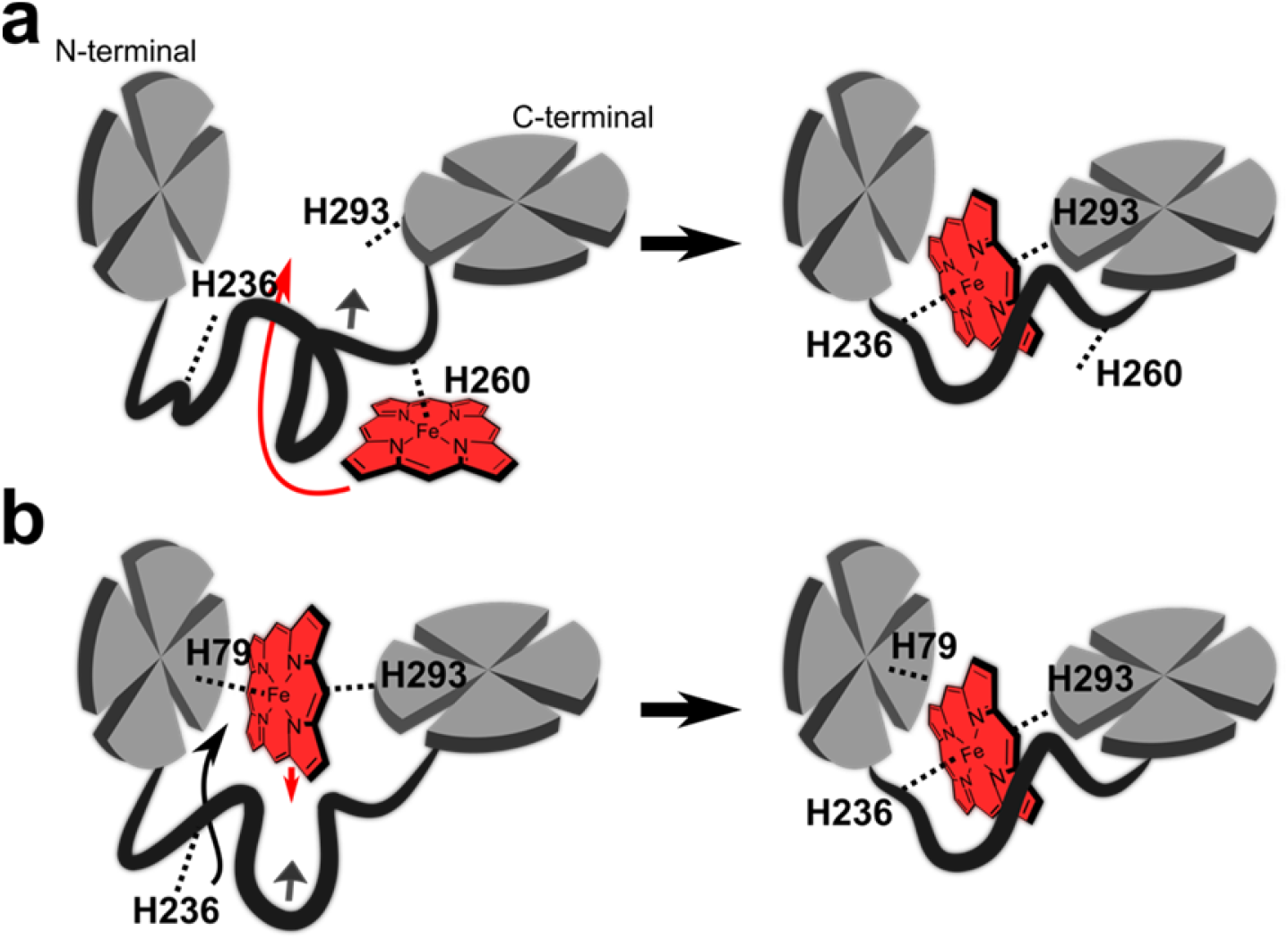
Possible heme-recruiting mechanisms involving novel heme-binding sites. It can be hypothesized that a heme-recruiting mechanism may be involved in guiding heme into its preferred binding pocket between H236 and H293. We propose the following mechanisms: **a** H260 initially binds heme on the surface of the protein, with H236 and H293 already in their correct positions to bind heme, a movement in the flexible loop causes H260 to shift into a less desirable position. This shift most likely propagates the transfer of heme from H260 into the binding pocket. **b** H79 binds heme in the known binding pocket, with H293 already in position. H236 is, however, not yet in a favourable position to form the H236/H293 binding complex. Movements in the flexible loop, shift H236 into an ideal binding position, with the position of H79 becoming unfavourable for heme binding. This leads to a downward movement of the heme into the binding pocket. After introduction of one heme molecule into the intended binding pocket, a second heme molecule can transiently bind to the other heme-binding motifs (e.g., H79, H105, H238, and H260) on the surface (not shown).

## Discussion

Hemopexin plays a major role in humans by detoxifying heme and preventing organismal damage under hemolytic conditions^5^. Several reports show its applicability as therapeutic agent^5,19,20,42,43^. This advancement in the medical application of hemopexin stands in strong contrast to the knowledge available regarding the exact nature of the heme-hemopexin interaction. To the best of our knowledge, the conflict between earlier studies and the crystal structure, which suggested different residues to be heme-coordinating, has not yet been resolved. Herein, we subjected long-standing assumptions of the heme-hemopexin interaction to thorough examination. We determined the heme binding to human hemopexin using SPR and were able to confirm high-affinity binding with a K_D_ value of 0.32 ± 0.04 nM for the first binding event. This value is higher than that reported by Hrkal *et al*. in 1974 (K_D_ value of <1 pM)^26^, but more in line with Morgan *et al*. 1972 (K_D_ value of <10 nM)^24^. The higher K_D_ value appears to be more realistic considering that hemopexin is believed to release heme under physiological conditions^5^. Our SPR as well as competition studies confirmed that hemopexin’s affinity for heme is higher than that of HSA *in vitro^5,6^*. This result is again in accordance with the physiological situation, where heme is assumed to be transferred from HSA to hemopexin^5^. Even though, the stoichiometry of the heme-hemopexin interaction has been reported to be 1:1 various studies also reported 2:1 (heme:hemopexin) ratios, as determined in the present study^23,24,26^. Taken together, our results clearly support the hypothesis of more than one heme-binding site in hemopexin.

The binding of two heme molecules to hemopexin can be explained by two possible scenarios (Fig. 4). Firstly, a recruiting mechanism delivering heme to the final binding pocket (H236 and H293) via H260 is conceivable. A movement in the flexible loop causes H260 to shift into a less desirable position and leads to the transfer of heme from H260 into the binding pocket (Fig. 4a). Secondly, H79, being located in the same binding pocket as H236 and H293, binds heme with H293 already in a favourable binding position. H236 is, however, not yet in form to complex with H293. A movement in the flexible loop, shifts H236, that is not yet in the position to form the H236/H293 binding complex, into an ideal binding position. This leads to a downward movement of the heme into the binding pocket to H236/H293 (Fig. 4b).

Secondly, an independent site away from the actual binding site exists within the protein. It can be hypothesized that the exposed sequence stretch around H260 (RCSP**H**LVLS, K_D_ = 0.16 ± 0.06 μM) interacts with HSA in a first step and takes over heme from HSA. However, in the crystal structure, heme binds to H236 and H293, which form a hexacoordinate complex within the binding pocket. The results of our spectroscopic analysis show that the isolated peptides H236 and H293 possess a relatively low affinity in relation to other hemopexin-derived peptides. It thus seems unlikely that heme is transferred directly to the pocket, when other high-affinity sites on the protein surface are exposed and available for binding. Intriguingly, H260 is located in close proximity to the binding pocket and could potentially direct heme into the pocket in the subsequent step. Interestingly, H238, another possible candidate within the recruiting mechanism, is found only two amino acids downstream of the confirmed binding site H236. Assuming the hypothesis of a second, independent binding site, heme could be bound axially between H105 and H150 as was suggested in earlier works^23,44^. Our experimental data, however, revealed that heme binding to H150 is hindered by the disulfide bridge between C149-C154 in the immediate proximity. In addition, this situation was not observed in our docking experiments, and it is thus unlikely that the orientation in the protein could still allow for binding.

It might be discussed that the heme-binding behaviour of hemopexin is altered by glycosylation^45,46^. However, N240 is the only glycosylation site close to a potential heme-binding site, i.e. H236 and H238. It thus seems unlikely that heme binding is hindered at H236 due to glycosylation as this residue was confirmed as heme binder in the crystal structure analysis.

In summary, we re-evaluate the heme-hemopexin interaction and propose a new K_D_ value together with a heme-recruiting mechanism. These novel insights may contribute valuable information regarding hemopexin’s function and applicability that are currently under investigation in other laboratories, such as its antithrombotic and antiinflammatory effect^20,43,47^. While hemopexin is currently being considered for the treatment of intracerebral hemorrhage (ICH)^48^, it might also be administered for the therapy of heme overload conditions, such as in sickle cell disease or β-thalassemia^47,49,50^. In these conditions, our results may be of special interest as hemopexin can be faced with an excess of heme, so that even less affine heme-binding sites may be utilized by the protein.

## Methods

### SPR binding studies

SPR measurements were performed three times in running buffer (20 mM Tris-HCl (pH 7.4), 150 mM NaCl, 0.05% polyoxyethylen(20)-sorbitan-monolaurate (Tween 20®) on a Biacore T200 instrument at 25 °C. Hemopexin isolated from human plasma (Athens Research & Technology, Athens, GA) was covalently immobilized on a CM5 sensor chip (GE Healthcare, Chicago, IL) by amine coupling. Purity of the protein was tested by SDS-PAGE and Coomassie staining. Hemopexin (pH 4.5, 10 μg/ml) was diluted in acetate buffer. Hemopexin was injected on an EDC/NHS activated flow cell at 10 μl/min flow rate until an immobilization level of 2116 RU was achieved. An activated/deactivated flow cell was utilized as reference. The affinity constant and kinetic information were generated via five consecutive injections of hemopexin with varying heme (Fe(III)protoporphyrin IX; Frontier Scientific, Logan, UT) concentrations (0.75 nM to 12 nM). The standard single-cycle kinetics method (Biacore T200 Control Software, GE Healthcare) was used. The flow rate for heme addition was 30 μl/min. Surface regeneration was performed by using 10 mM NaOH. The resulting data were fitted with Biacore T200 Control Software (GE Healthcare) using a kinetic global fit 1:1 binding model, since the first, high-affinity heme-binding interaction was found to be 1:1.

### Homology modelling of hemopexin

A three-dimensional model of hemopexin’s structure was derived through homology modelling done via YASARA version 19.9.17^33^. The template search was done by running three iterations of Position-Specific Iterative - Basic Local Alignment Search Tool (PSI-BLAST), with an E-value threshold of 0.1, against UniRef90^51^ to build a position-specific scoring matrix (PSSM) from related sequences. By using this profile, the Protein Data Bank (PDB)^52^ was searched for potential modelling templates. False positives were avoided through the exclusion of common protein purification tags. Templates were ranked according to the BLAST alignment score, as well as the WHAT_CHECK^53^ score of structural quality that was obtained from the PDBFinder2 database^54^. The target sequence was fed to the PSI-Pred server for secondary structure prediction^55^. The monomeric model was built on the 1QJS-A-template^23^, which was regarded as the only suitable template for the given target sequence. Fifty conformations were sampled to model the loop regions. The quality of the final homology model assessed via the overall z-score, calculated as the sum of the individual dihedral, 1D packing and 3D packing z-scores. A pairwise structure-based MUSTANG alignment^56^ was conducted with YASARA to analyse the structural similarity between the PDB ID: 1QJS crystal structure^23^ and the homology model.

### Molecular dynamics (MD) simulations of the hemopexin homology model

The homology model was used without any adaptions as the holo-hemopexin structure. In order to create an apo-hemopexin model, which was used to investigate the hypothesized recruiting mechanism of heme by hemopexin, heme was removed from the holo-hemopexin homology model. Both the apo- and holo-hemopexin models were subjected to a 200 ns long, explicit, all-atom MD simulation in solution conducted in YASARA 19.9.17^33^. The purpose of this simulation was to generate a conformational ensemble of the protein models providing insights into their structural dynamics, and to ensure their conformational stability. YASARA enabled automatic parameterization of the heme-protein complex with the force field (AMBER11^40^) using the AutoSMILES method as done in previous studies^33,57^. The simulation setup included a simulation cell with boundaries extending at least 10 Å on each side from all protein molecules, optimization of the hydrogen bond network^58^, as well as a pK_a_ prediction to refine the protonation states of the protein residues at a pH of 7.4^58^. Sodium or chloride ions were added to replace water molecules in the simulation system to achieve a physiological concentration of 0.9%. Steepest descents and simulated annealing minimizations were carried out to remove clashes. These minimizations were followed by a simulation for 200 ns using the AMBER11 force field^40^. Periodic boundary conditions were used with an 8 Å cut-off for long range forces. Long range electrostatics were accounted for using the particle mesh Ewald method^59^. Simulation snapshots were written to disk every 200 ps. The analysis tools available in YASARA^33^ and VMD 1.9.3^60^ were used to compute MD-derived observables of structural and conformational change. Molecular graphics were created using YASARA^33^ and VMD 1.9.3^60^. Plots were created using the Grace 5.1.25 program^61^.

### Qualitative determination of disulfide bonds in hemopexin

In order to confirm that all cysteine residues of hemopexin are involved in disulfide bonds, 2-iodoacetamide (4 mM) in 10 mM phosphate buffer (pH 7.4) was added to a solution of hemopexin in PBS buffer (pH 7.4). The molar mass of the derivatized protein was compared to the wild-type untreated hemopexin as determined by MALDI mass spectrometry using an ultrafleXtreme TOF/TOF mass spectrometer (Bruker Daltonics, Billerica, MA) with 2,5-dihydroxyacetophenone (Fluka, Buchs, Switzerland) as matrix. Additionally, Ellman’s test^35^ was performed with hemopexin in comparison to cysteine (positive control) and cystine (negative control) by mixing 150 μl of the sample solutions with 100 μl Ellman’s reagent (1.76 mg/ml 5,5’-dithiobis-(2-nitrobenzoic acid) in 0.2 M sodium phosphate buffer, pH 8). Absorption was measured after 10 min incubation at 410 nm.

### Peptide synthesis

Standard Fmoc solid phase peptide synthesis was used to produce the peptides as described earlier^37^. Crude peptides were purified after cleavage from the resin via semipreparative HPLC and quantified via amino acid analysis as described earlier^37^. The intramolecular disulfide bond of peptide H150 (AVECHRGEC) was formed by air oxidation using a previously established protocol^62^. Complete oxidation was confirmed using Ellman’s test as described above^35^.

### Mass spectrometry

The molar masses of the synthesized peptides and the modified protein were determined using a MALDI TOF/TOF mass spectrometer ultrafleXtreme (Bruker Daltonics GmbH). The matrix used was α-cyano-4-hydroxycinnamic acid (Sigma Aldrich, St. Luis, MO).

### UV/Vis spectroscopy

Analysis of heme binding for K_D_ determination was performed by UV/Vis spectroscopy on a Multiskan GO microplate spectrophotometer (Thermo Fisher Scientific, Waltham, MA) as described earlier^37^. The peptide concentration was 20 μM in 100 mM *N*-2-hydroxyethylpiperazine-*N*’-2-ethanesulfonic acid (HEPES) (Sigma Aldrich) buffer (pH 7.4). Heme stock solutions were prepared at 1 mM concentration as described earlier^37^, and freshly diluted in buffer immediately before usage. Heme concentrations were normalized using an extinction coefficient of 32.482 mM^-1^cm^-1^ (398 nm) in HEPES^32^. Evaluation of the UV/Vis spectra was performed as described earlier using PRISM 7, GraphPad Software^37^.

### rRaman spectroscopy

The Resonance Raman spectra were collected with a commercial micro-Raman set-up (CRM 300, WITec GmbH, Germany) with 405 nm diode laser as excitation wavelength. The spectrometer is fitted with a grating of 1800 g/mm and a CCD camera (DV401A BV-532, ANDOR, 1024 x 127 pixels). The samples were measured in liquid condition using Zeiss 10x objective (NA 0.2) with a laser power of 22 mW before objective. The sample was taken in a quartz cuvette (~80 μwww of sample volume) and placed on home-built sample holder which was spirally rotated with a speed of 1 rotation/60s. Raman spectra were collected in triplicate with 30s integration time per spectrum. Resonance Raman spectra were background corrected using SNIP algorithm^63^ and mean spectra were generated. For analysis GNU R platform^64^ and in-house built script was used. For display the spectral region of interest (600 cm^-1^ to 900 cm^-1^ and 1400 cm^-1^ to 1800 cm^-1^) was plotted.

### Competition assay

Heme was dissolved in 30 mM NaOH for 30 min at 25 °C in the dark. The solution was diluted to 100 μM in PBS buffer (pH 7.4) (stock solution). The peptides H105, H236, H238, H260, and H293 were dissolved in PBS buffer (pH 7.4) to a final concentration of 100 μM (stock solutions). Heme-peptide complexes (1:1) were prepared from the stock solutions, diluted to 20 μM (PBS), and incubated for 30 min before measurement. To 100 μl of the complexes (20 μM) either 100 μl of a hemopexin solution (10 μM) or buffer was added. The samples were then transferred to high precision quartz micro cuvettes (Hellma Analytics, Müllheim, Germany). Two sets of spectra ranging from 230 to 750 nm were recorded simultaneously using two UV-3100PC UV/Vis spectrophotometers (VWR, Radnor, PA). In the first device, the heme-peptide complex incubated with hemopexin was measured, whereas in the second instrument the control (heme-peptide complex only) was recorded. Spectra were measured every 95 s for 4 h at 25 °C. Data analysis was performed with PRISM 7 (GraphPad Software, San Diego, CA). For the measurements of the heme-HSA complex, a HSA solution (100 μM stock) was used instead of peptide in the same way as described above.

### Molecular docking of heme onto the apo- and holo-hemopexin models

Both the apo- and holo-hemopexin models, were subjected to molecular docking studies. This was done in order to validate heme binding to the experimentally established heme-binding motifs, as well as to gain insight into the binding. The ChemSpider database^65^ was used to retrieve the structure of the heme ligand (ChemSpider ID 16739951). In order to improve efficiency of the docking process, the docking search space was restricted to a 15 Å x 15 Å x 15 Å cubic simulation cell around a single experimentally determined nonapeptide HBM. An “ensemble docking” approach^66^, based on the AutoDock Vina algorithm^67^ was used to execute the docking process via YASARA. The ensemble docking approach produced a receptor ensemble consisting of 10 high-scoring sidechain rotamers solutions thereby introducing receptor flexibility and improving the docking process as opposed to a fully rigid receptor. The heme was docked 50 times to each member of the ensemble, resulting in 500 docking runs per HBM. The docking results were clustered based on predicted binding energies by the Vina scoring function and a 5 Å RMSD threshold between different poses of the heme was used as criterion for specific cluster membership. This process was repeated for each of the experimentally determined nonapeptide HBMs.

### Molecular dynamics simulations of the heme-bound complexes

The stability of docked receptor-ligand complexes in solution cannot be inferred or validated from the docking results alone^68^. In order to validate the docking process, a selected docked complex of each HBM, for both the apo- and holo-hemopexin, was subjected to a 500 ps refinement simulation via YASARA^41^. This refinement simulation used the YAMBER3 force field^69^. A snapshot was saved every 25 ps and the energy was minimized^69^. The purpose of this refinement simulation was to validate the newly created receptor-ligand complex and to identify the structure with the lowest energy. This lowest energy structure from the refinement simulations were further subjected to two independent 20 ns MD simulations using an identical simulation setup. Additionally, an estimation binding energy analysis of each simulated complex, was done via YASARA. The program approximates the binding energy (kJ*mol^-1^) by applying the following formula: Binding Energy= E_pot Receptor_ + E_solv Receptor_ + E_pot Ligand_ + E_solv Ligand_ - E_pot Complex_ - E_solvComplex_, with E_pot_ being the potential energy and Esolv the solvation energy. The calculated binding energies were obtained from the MD simulations of the docked complexes. This estimation of binding energy in an explicit MD system in solution, resulted in relevant docking energies, unlike the Autodock Vina *in vacuum* binding energy calculations. Since the calculated binding energy results are only approximations, the results can only be used to assess the quality of binding between different motifs within a particular protein structure.

## Data Availability

The datasets generated and analysed during the current study are available from the corresponding author D.I. on reasonable request.

## Acknowledgements

The authors would like to thank Sabrina Linden (University of Bonn, Germany) for technical assistance. Financial support by the University of Bonn is gratefully acknowledged.

## Author Contributions

D.I. designed and planned the project. M.S.D, B.F.S., and D.I. conceived the study and workflow. M.S.D., M.-T.H., B.F.S., and A.R. performed the experiments and collected the data. M.S.D., B.F.S., A.A.P.G., M.-T.H., A.R., U.N., and D.I. analysed the data. A.A.P.G. and F.S. performed the computational studies. The manuscript was written through the contribution of all authors.

## Competing interests

The authors declare no competing interests.

## Materials & Correspondence

Correspondence and requests for materials should be addressed to D.I.

## Notes

### Competing Interest Statement

The authors have declared no competing interest.

