## Supplementary Information for "Revisiting the interaction of heme with hemopexin: Recommendations for the responsible use of an emerging drug"

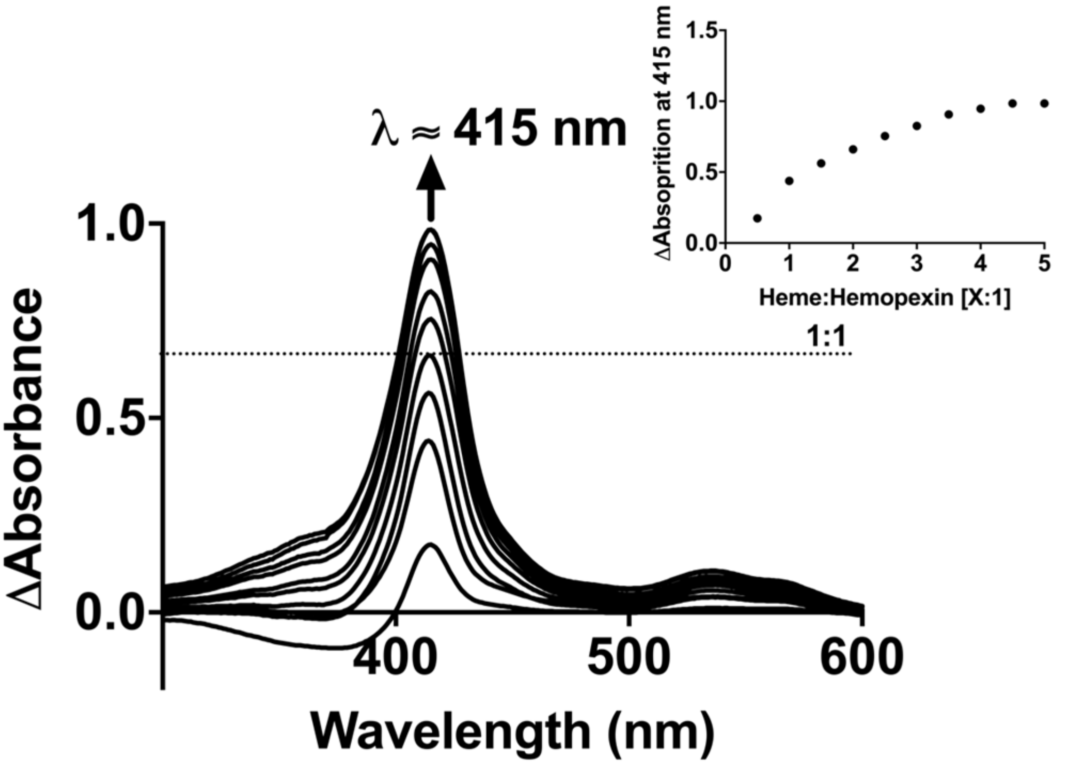


**Fig. 1**. **Titration of hemopexin with heme shows an increase in absorbance.**

Difference spectra of the heme-hemopexin complex obtained from the interaction of hemopexin (5 μM) with increasing heme concentrations (2.5 – 23.8 μM). Notably, the resulting curve (inset) is not characteristic of a single high-affinity binding site. The absorbance at the Soret band increases at heme:hemopexin ratios exceeding 1:1.

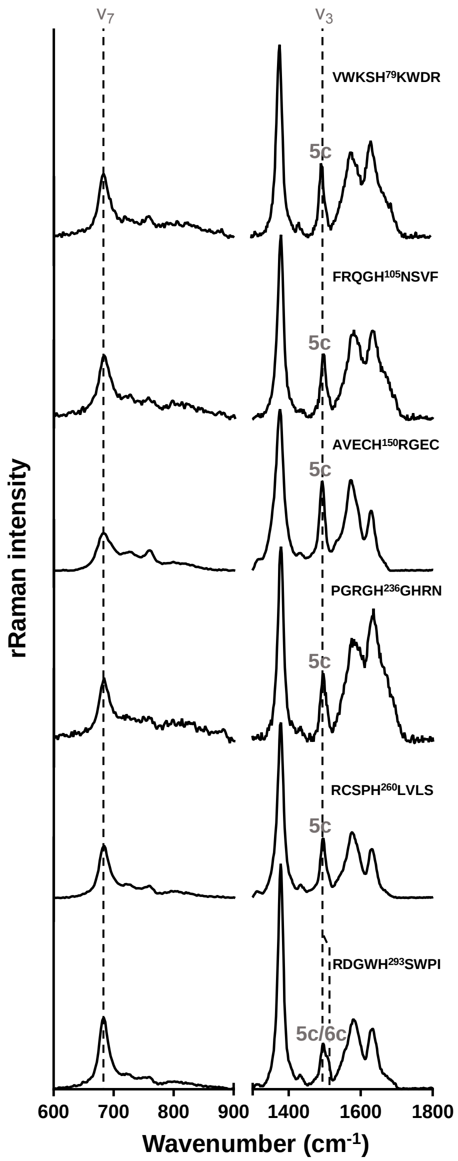


**Fig. 2. Resonance Raman spectra of heme-peptide complexe**s.

All complexes were measured at 400 µM in PBS with excitation at 405 nm. As indicated by the $\nu_3$-vibrational band, all peptides bound heme in a pentacoordinated fashion, with the exception of peptide motif H293, which showed a mixture of penta- as well as hexacoordination. Peptide H238 could not be measured due to insolubility of the complex.

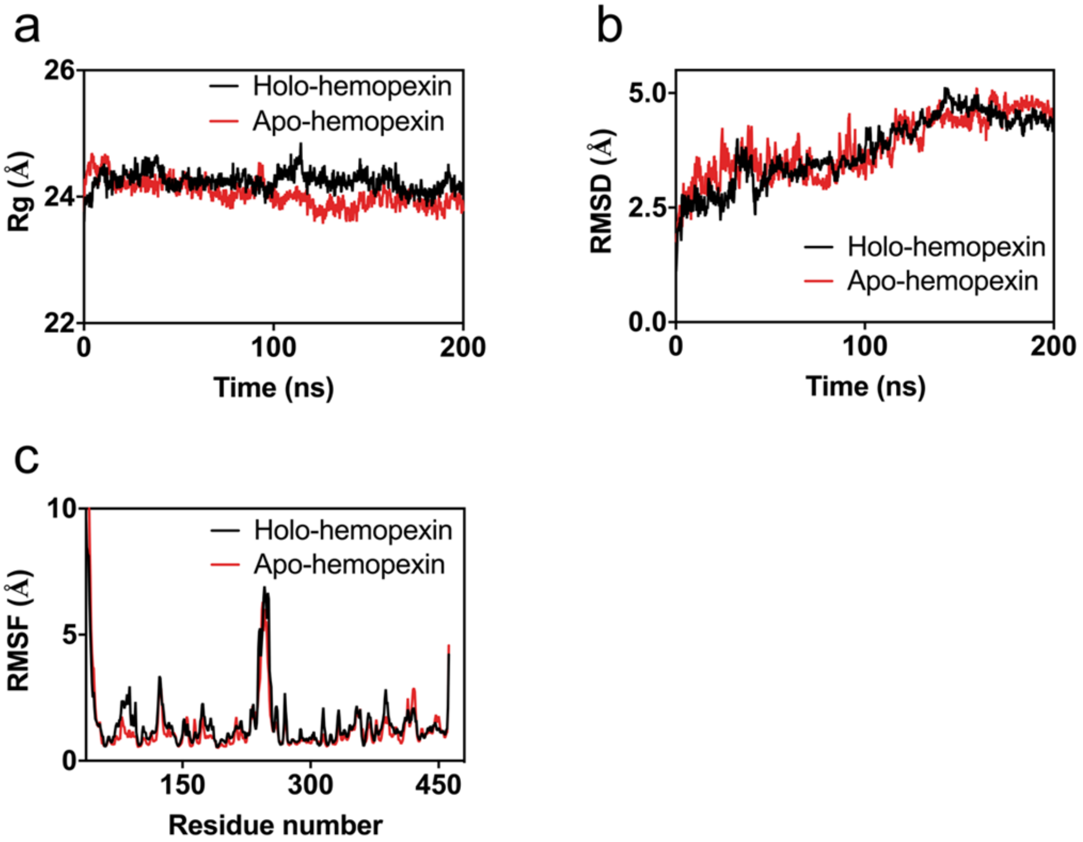


**Fig. 3. The root-mean-square deviation (RMSD), root-mean-square fluctuation (RMSF), and radius of gyration (Rg) profiles of the apo- and holo-hemopexin models determined from 200 ns long MD simulations.**

**a** The backbone RMSD value of apo-hemopexin was calculated as 3.84 ± 0.67 Å with a range of 1.70 Å to 5.16 Å. The backbone RMSD value of holo-hemopexin was calculated as 3.76 ± 0.75 Å with a range of 1.05 Å to 5.27 Å. During both simulations the RMSD values started to stabilize after approximately 170 ns, indicating a stable structure. **b** The Rg values for both, the apo-hemopexin and holo-hemopexin, did not show any significant fluctuations. These slight fluctuations in the overall radius of gyration indicate structural stability during both simulations. Unfolding of the protein structure did not occur. **c** The RMSF values were calculated per residue to elucidate on whether the predictive binding sites do have an influence on the protein's conformation. As expected, significant fluctuations occur at the N-terminal of the protein and to a lesser extent at the C-terminal. The peak patterns between the apo-hemopexin and the holo-hemopexin are similar. This may indicate that having a single heme in the structure does not have a significant impact on either the other heme-binding motifs or on the conformational changes the protein undergoes.

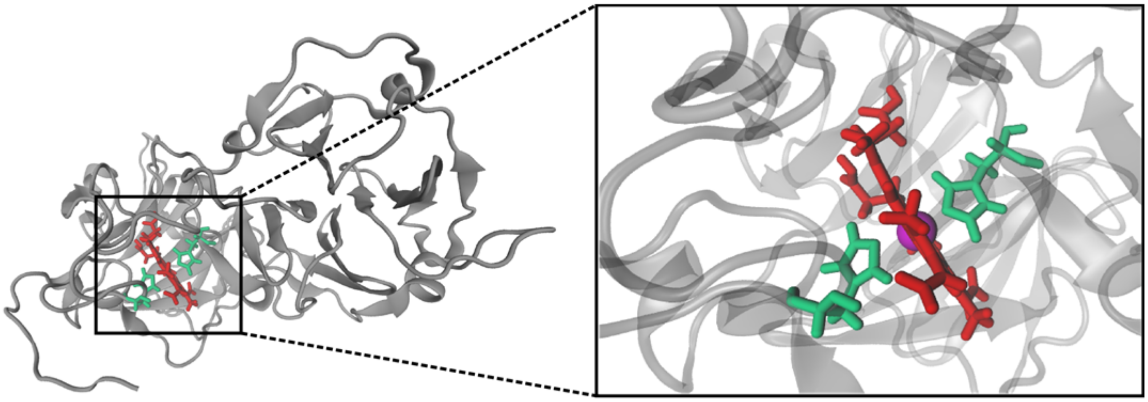

**Fig. 4. The homology model of human hemopexin accommodates heme in the confirmed binding pocket.**

Validation of the homology model (grey ribbons) by docking of heme. Heme (red sticks) docks into the known H236/293 (green sticks) binding pocket observed in the crystal structure, a close-up view of the complex is shown on the right.

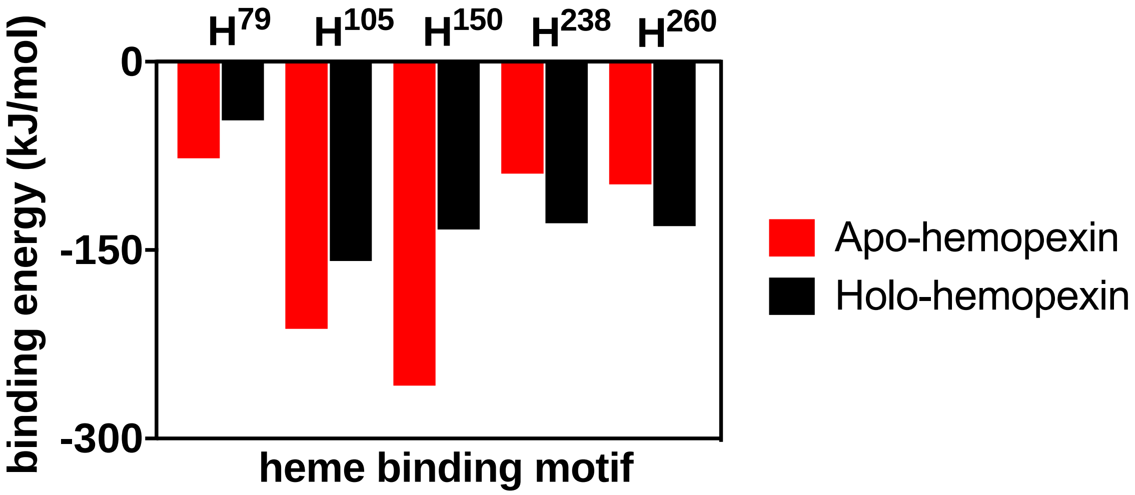


**Fig. 5. The average binding energy (in kJ/mol) of potential coordinating histidines in both apo- and holo-hemopexin docking complexes.**

The binding energy values are obtained from explicit 20 ns MD simulations of the docked complexes in solution that were formed when heme was docked on the specific heme-binding motif of hemopexin. red bars represent apo-hemopexin, black bars represent holo-hemopexin. Heme-binding motifs with higher binding energy values (H79, H238 and H260) indicate stronger binding of heme.

| Sequence | $K_D$ [µM] | $\lambda$ [nm] | Posttranslational modification | Disulfide bond | SeqD-HBM | Reference |
|---|---|---|---|---|---|---|
| VWKS**H**$^{79}$KWDR | 3.72 ± 0.23 | 420 | - | - | -* | [1,2] |
| FRQG**H**$^{105}$NSVF | 0.48 ± 0.24 | 419 | - | - | + | [3] |
| AVEC**H**$^{150}$RGEC | n. b. | - | - | C154 | -** | [1–3] |
| HRGE**C**$^{154}$QAEG | n. b. | - | - | C154 | + | |
| PGRG**H**$^{236}$GHRN | 3.61 ± 0.37 | 421 | N-linked at N240 | - | + | [3] |
| RGHG**H**$^{238}$RNGT | 0.35 ± 0.17 | 420 | N-linked at N240 | - | + | |
| GNST**H**$^{249}$HGPE | n. b. | - | N-linked at N246 | - | + | |
| NSTH**H**$^{250}$GPEY | n. b. | 421 | N-linked at N246 | - | + | |
| HGPE**Y**$^{254}$MRCS | n. b. | - | - | C257 | + | |
| RCSP**H**$^{260}$LVLS | 0.16 ± 0.06 | 420 | - | C257 | + | |
| RDGW**H**$^{293}$SWPI | 5.67 ± 0.58 | 420 | - | - | + | [3] |
| TKGG**Y**$^{335}$TLVS | n. b. | - | - | - | + | |
| VTSLLGCT**H**$^{462}$ | n. b. | - | - | C460 | + | |

**Table 1. Summary of the synthesized peptides**.

For the peptides in complex with heme the following properties are given: $K_D$ values, shift of the absorption maximum ($\lambda$), posttranslational modifications[4] and disulfide bonds[5] occurring in the protein, prediction by the algorithm SeqD-HBM and reference, where applicable. *This motif was excluded by WESA. **This motif was excluded due to an intramolecular disulfide bond.
